## Supplementary figures 1 to 7 and tables S1 and S2 for "The structure and flexibility analysis of the *Arabidopsis* Synaptotagmin 1 reveal the basis of its regulation at membrane contact sites"

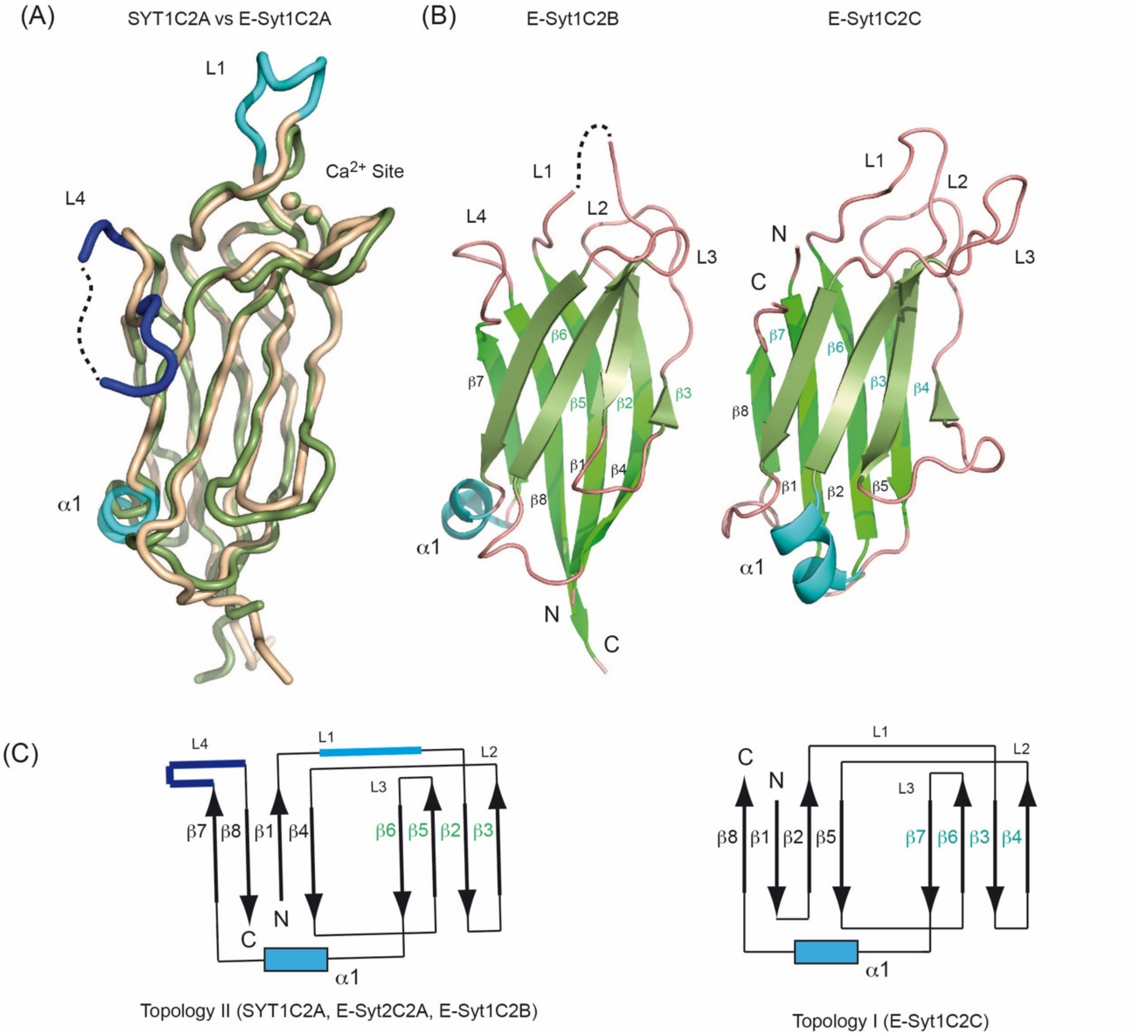


**Figure S1.** (A) Superposition of the structures of SYT1C2A (green) and E-SYT1C2A (wheat) (B) A ribbon representation of the structure of E-Syt2C2B (left) and E-Syt2C2C (right). (C) Schematic representation of the topologies of the C2 domains.

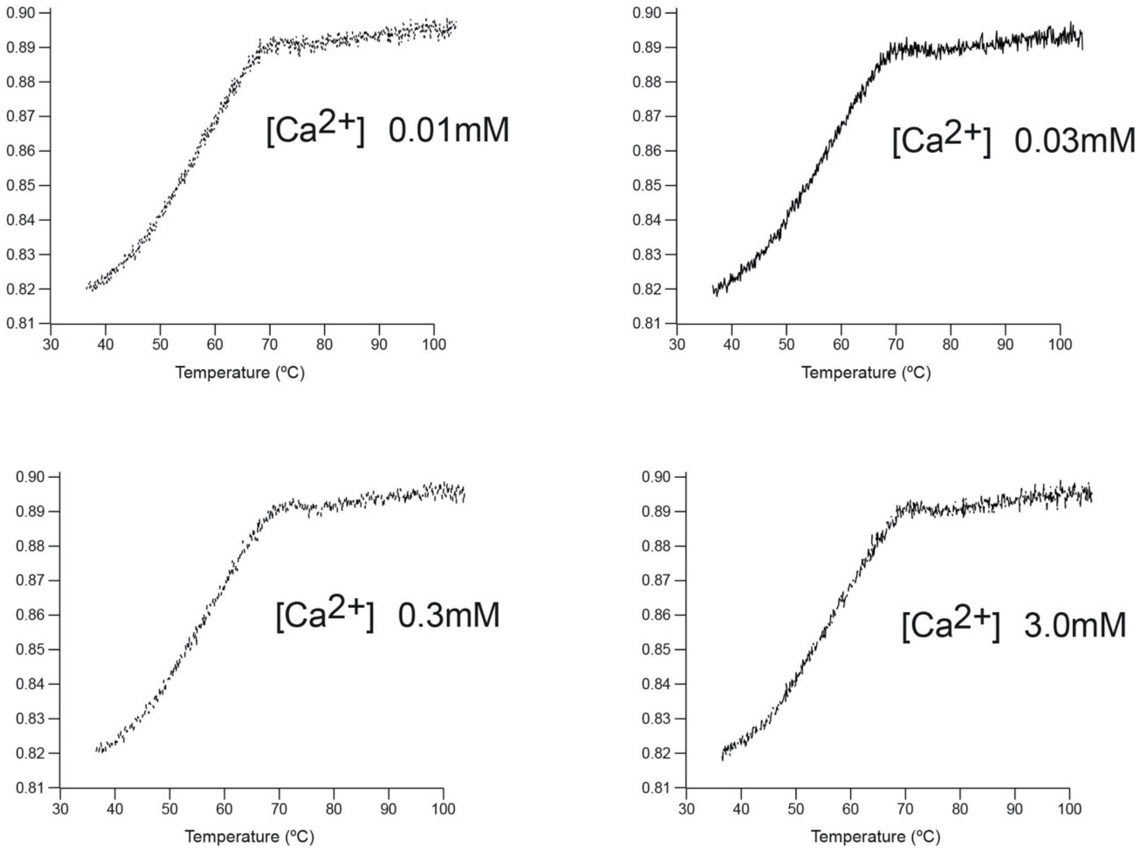


**Figure S2.** Thermal denaturation of SYT1C2B at different $Ca^{2+}$ concentrations. The sequence analysis of SYT1C2B shows that the protein fragment does not harbor any of the residues conforming the $Ca^{2+}$ binding site (figure 1C). Thus, as expected, no shift in the Ti nor in the initial fluorescence ratio is observed upon $Ca^{2+}$ addition, proving that the protein fragment does not bind $Ca^{2+}$.

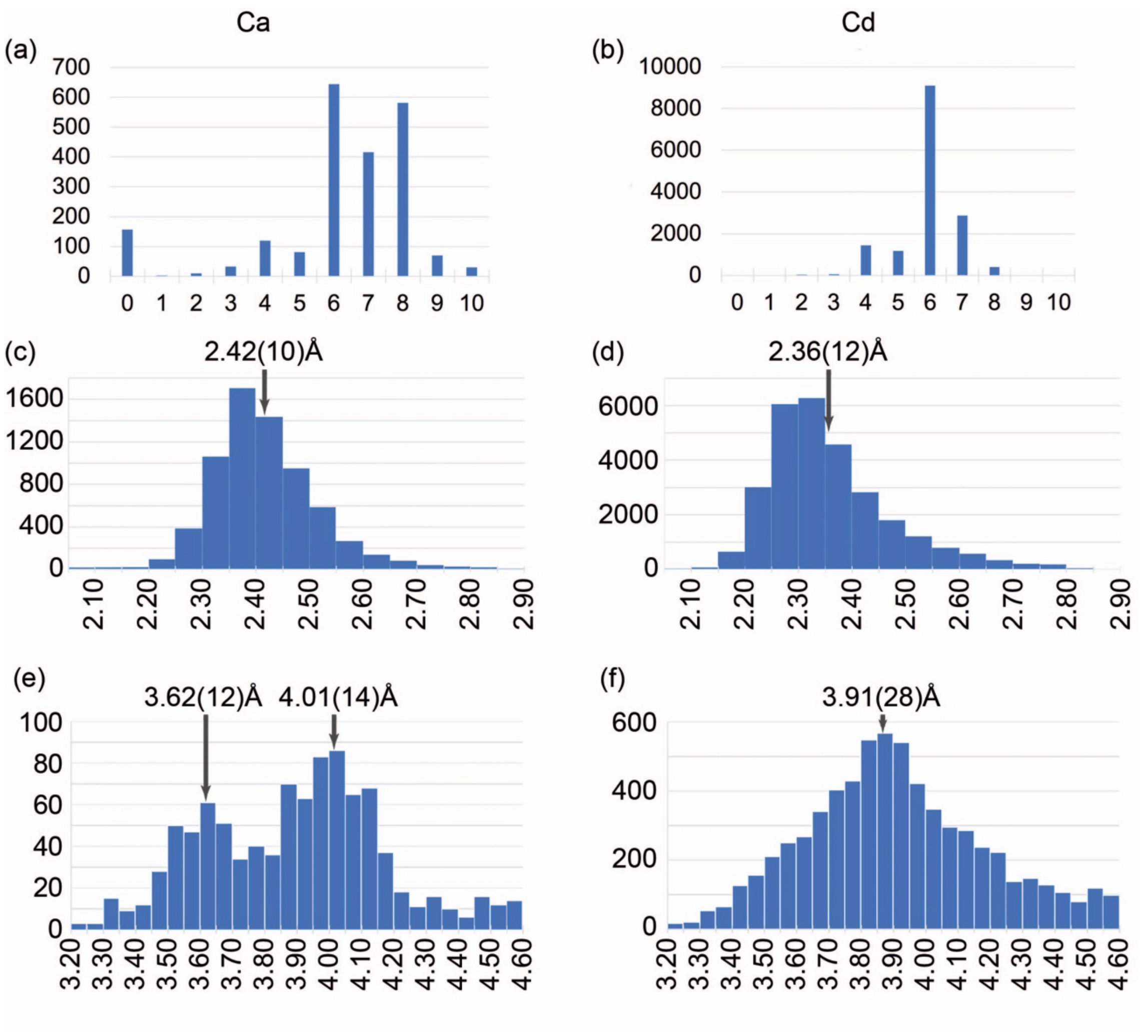


**Comparison of Ca (left column) and Cd (right column) environment in the Cambridge Structural Database (CSD).** The more likely geometry corresponds to the more stable structures. (a, b) Distribution of the number of connections for Ca and Cd atoms in the CSD. (c, d) Distribution of Ca/Cd-O distances (Å). (e, f) Observed Ca-Ca and Cd-Cd distances (Å) The mean and the standard deviations of the maximum are given in the histograms.The metal binding site of SYT1C2A restricts the metal-metal distance to 3.7A. This is in agreement with the first maximum observed for Ca-Ca distribution (panel e). Conversely, the Cd-Cd distribution displays a maximum at 3.9A (panel f). Hence, it is likely that the constraints imposed at SYT1C2A site hinders the presence of two Cd atoms.

**Figure S3.** Analysis of the coordination and geometry of the interaction between $Ca^{2+}$, $Cd^{2+}$ and oxygen using crystallographic data from the CSD.

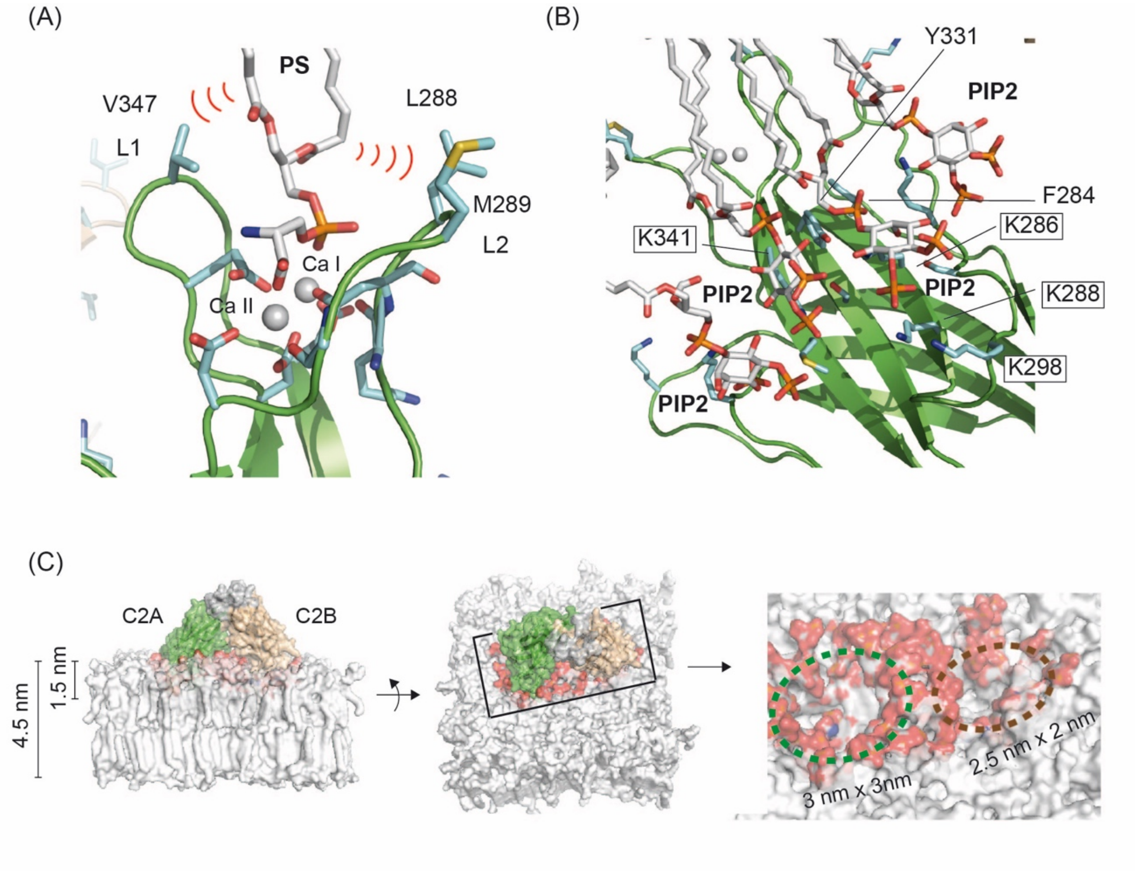


**Figure S4.** Two details of the MD simulations showing the relevant interactions of SYT1C2A and PSPI membrane at the $Ca^{2+}$ (A) and polybasic (B) lipid binding sites. Those residues mutated to generate the SYT1C2A-PolyB fragment are highlighted (C) The insertion of the SYT1C2AB tandem distorts membrane structure by defining large cavities into one leaflet. The bilayer is depicted as a white surface and the lipids interacting with the C2AB tandem are displayed in red.

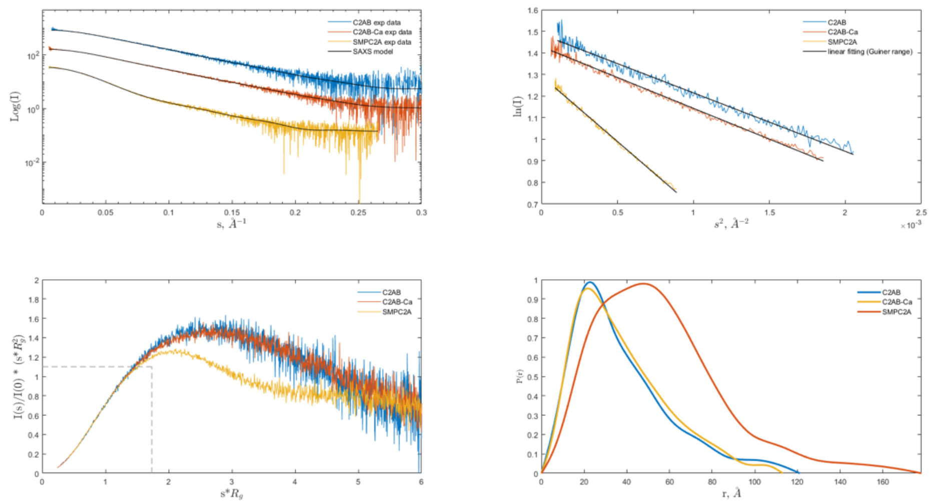


**Figure S5.** SAXS data and their analysis for C2AB, C2AB-Ca and SMPC2A (A) Experimental scattering curves compared to the fit (black solid traces) of the simulated curves calculated from MultiFoxs modelling (the $\chi 2$ values are reported in the Figure S6) (B) Guiner analysis. The black solid trace represents the linear fit from which $R_g$ was calculated. (C) Dimensionless Kratky plot. The intersection of the dotted black trace corresponds to the value for bovine serum albumin. (D) Pair-distance distribution function plot, P(r).

C2AB

Starting model: Rg= 25.82 Dmax= 87.97 $\chi^2$= 2.00

| State | Conformation | fraction | Rg | Dmax | $\chi^2$ |
|---|---|---|---|---|---|
| 1 | 1 | 1.00 | 26.87 | 99.31 | 1.20 |
| 2 | 1 | 0.54 | 22.85 | 77.53 | 1.10 |
| | 2 | 0.46 | 34.10 | 113.80 | |
| 3 | 1 | 0.39 | 22.26 | 78.93 | 1.07 |
| | 2 | 0.14 | 24.45 | 83.92 | |
| | 3 | 0.47 | 34.10 | 113.80 | |
| 4 | 1 | 0.24 | 21.26 | 74.33 | 1.07 |
| | 2 | 0.11 | 23.68 | 84.51 | |
| | 3 | 0.19 | 24.45 | 83.92 | |
| | 4 | 0.47 | 34.10 | 113.80 | |

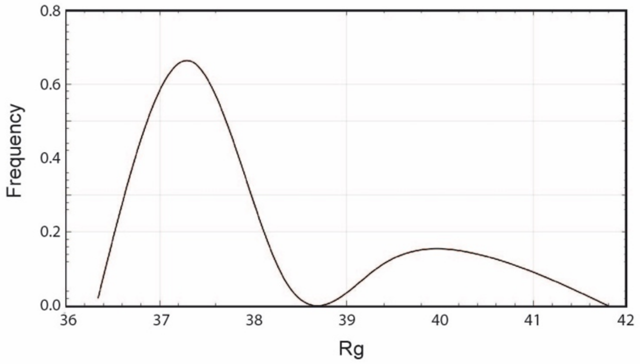


C2AB-Ca

Starting model: Rg= 25.82 Dmax= 87.97 $\chi^2$= 2.99

| State | Conformation | fraction | Rg | Dmax | $\chi^2$ |
|---|---|---|---|---|---|
| 1 | 1 | 1.00 | 28.06 | 102.30 | 1.27 |
| 2 | 1 | 0.53 | 23.45 | 81.01 | 1.06 |
| | 2 | 0.47 | 34.83 | 116.87 | |
| 3 | 1 | 0.24 | 21.99 | 71.93 | 1.04 |
| | 2 | 0.30 | 25.32 | 84.81 | |
| | 3 | 0.46 | 34.83 | 116.70 | |
| 4 | 1 | 0.10 | 21.17 | 74.74 | 1.04 |
| | 2 | 0.16 | 22.66 | 76.32 | |
| | 3 | 0.29 | 25.32 | 83.92 | |
| | 4 | 0.46 | 34.83 | 116.70 | |

C2AB

Starting model: Rg= 25.82 Dmax= 87.97 $\chi^2$= 2.00

| State | Conformation | fraction | Rg | Dmax | $\chi^2$ |
|---|---|---|---|---|---|
| 1 | 1 | 1.00 | 26.87 | 99.31 | 1.20 |
| 2 | 1 | 0.54 | 22.85 | 77.53 | 1.10 |
| | 2 | 0.46 | 34.10 | 113.80 | |
| 3 | 1 | 0.39 | 22.26 | 78.93 | 1.07 |
| | 2 | 0.14 | 24.45 | 83.92 | |
| | 3 | 0.47 | 34.10 | 113.80 | |
| 4 | 1 | 0.24 | 21.26 | 74.33 | 1.07 |
| | 2 | 0.11 | 23.68 | 84.51 | |
| | 3 | 0.19 | 24.45 | 83.92 | |
| | 4 | 0.47 | 34.10 | 113.80 | |

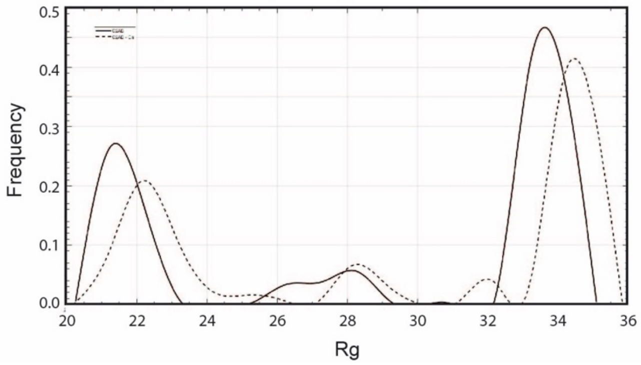


**Figure S6.** MultiFoXS output for SMPC2A, C2AB-Ca and C2AB. The tables show the top scoring solutions for several state models together with the Rg distribution for the conformations generated by the server. More than 10,000 conformations were calculated. The histograms display two obvious relative modes that correspond to two different conformations. Note the large difference between these modes for the C2AB fragment

```
> SYT1   1 MGFFSTILGFCGFGVGISLGLVIGYVLFVYLLPN 36
> SYT3   1 MGFFTSVLGIIGFVIGIPIGLILGFFVLIYSQPS 36
PSIPRED     HHHHHHHHHHHHHHHHHHHHHHHHHHHHHHHH
SPIDER3     HHHHHHHHHHHH HHHHHHHHHHHHHHHHHHH
PSSPRED4    HHHHHHHHHHHH      HHHHHHHHHHHHH
DEEPCNF     HHHHHHHHHH HHHHHHHHHHHHHHHHHHHHH

> E-Syt2   ARSFALLLPVYALGYLGLSFSWVLLALALLAWCRR
> Tcb1     SVAIFFIGGVASFALGHYKFSMGSAFFVIVITSLLYRT

> PDB1WAZ  PVLVILLGVVGLSALTGYLDYVLLPALAIFIGLTIYAIQR
  SS 1WAZ   HHHHHHHHHHHHHH         HHHHHHHHHHHHHH
```

Quick2D gives you an overview of secondary structure features like alpha-helices, extended beta-sheets, coiled coils, transmembrane helices and disorder regions.

**Figure S7**. Comparison of the hydrophobic regions of plant SYTs, human E-Syt1, Trcb1 and a buried helical hairpin with known structure. The secondary structure, as predicted for SYTs with various algorithms, is also displayed (Quik2D from the HHpred server).

**Table S1. Data collection and refinement statistics**

| | | |
|---|---|---|
| **Data collection** | | |
| Construct | SYT1C2A Ca | SYT1C2A Cd |
| Space group | R 3 :H | R 3 :H |
| a, b, c, Å | 69.13 69.13 115.67 | 70.94 70.94 116.31 |
| α, β, γ, ° | 90 90 120 | 90 90 120 |
| Resolution, Å | 41.59-2.10 (2.16-2.10) | 42.23-2.00 (2.06-2.00) |
| Rpim | 0.023 (0.706) | 0.053 (1.041) |
| $CC_{1/2}$ | 0.999 (0.551) | 0.997 (0.449) |
| I/σ(I) | 14.3 (1.1) | 7.6 (0.8) |
| Completeness, % | 100.0 (100.0) | 99.5 (99.1) |
| Redundancy | 6.6 (6.8) | 3.4 (3.5) |
| **Refinement** | | |
| Resolution, Å | 41.59-2.10 (2.18-2.10) | 42.23-2.00 (2.16-2.00) |
| No. of reflections | 24010 | 14573 |
| Rwork/Rfree | 20.7/26.9 (35.8/38.9) | 18.6/21.1 (34.3/35.1) |
| No. of atoms | | |
| Protein | 1054 | 1074 |
| Calcium/Cadmium ions | 2 | 2 |
| Zinc/Nickel ions | 1 | 2 |
| Chlorine ions | 1 | 1 |
| Ethylene glycol | - | 2 |
| Water molecules | 10 | 28 |
| Average B factors | 92.23 | 62.98 |
| rmsd | | |
| Bond lengths, Å | 0.009 | 0.008 |
| Bond angles, º | 0.971 | 0.917 |
| Ramachandram plot | 96.0% in the core<br>4.0% in the allowed | 96.85% in the core<br>3.15% in the allowed |

Highest-resolution shell is shown in parenthesis

**Table S2.** Data collection and structural parameters by SAXS.

| | C2AB | C2AB-Ca | SMPC2A(N34-E397) |
|---|---|---|---|
| **Data collection parameters** | | | |
| BeamLine | B21, Diamond Light Source, Harwell (UK) | | |
| Detector | Eiger 4M | Eiger 4M | Pilatus 2M |
| Beam size (mm) | 0.34 x 0.40 | 0.34 x 0.40 | 0.2 x 0.2 |
| Energy (keV) | 12.4 | 12.4 | 12.4 |
| Sample-to-detector distance (mm) | 3700 | 3700 | 4014 |
| q range (A-1) | 0.0026 – 0.34 | 0.0026 – 0.34 | 0.0038 - 0.42 |
| Exposure time (s) | 3 | | |
| Temperature (K) | 293 | | |
| Data collection mode | SEC online | | |
| **Structural parameters** | | | |
| Concentration range (mg ml$^{-1}$) | 10 | 10 | 6.6 |
| $q$ Interval for Fourier inversion (Å$^{-1}$) | 0.011 – 0.258 | 0.007 – 0.258 | 0.010 -0.190 |
| $R_g$ [from P(r)] (Å) | 32.1 | 30.6 | 45.1 |
| $R_g$ [from Guiner approximation] (Å) | 28.2 | 29.1 | 44.9 |
| $sR_g$ limits [from Guiner approximation] | 0.33 1.30 | 0.21 1.30 | 0.41 1.28 |
| Dmax (Å) | 121 | 114 | 176 |
| Porod coefficient | 2.0 | 2.1 | 2.8 |
| Porod volume estimate (nm$^3$) | 50 | 54 | 138 |
| DAMMIF excluded volume (nm$^3$) | 55 | 57 | 151 |
| Molecular Mass (kDa) | | | |
| From Porod volume (x0.53) | 28 | 29 | 73 |
| From excluded volume (x 0.5) | 28 | 29 | 76 |
| From sequence | 33 | 33 | 41 per protomer |
| Modelling | | | |
| Ambiguity | 1.0 (is unique) | 1.5 (might be ambiguous) | 2.1 (might be ambiguous) |
| Resolution (Å) | 30 ± 2 | 31 ± 3 | 57 ± 4 |
| SASBDB | SASDKJ9 | SASDKK6 | SASDKG6 |
| **Software employed** | | | |
| Primary data reduction | DAWN pipeline (Diamond Light Source, UK) | | |
| Data processing | ScÅtter IV | | |
| Computation of model intensities | CRYSOL | | |
| Flexibility | MultiFoxs | | |

$q = 4\pi sin(\theta)/\lambda$, where $2\theta$ is the scattering angle and $\lambda$ is the wavelength od incident X ray beam
