## Supplementary figures and images for "The structure and flexibility analysis of the *Arabidopsis* Synaptotagmin 1 reveal the basis of its regulation at membrane contact sites"

### Supplementary movie 1

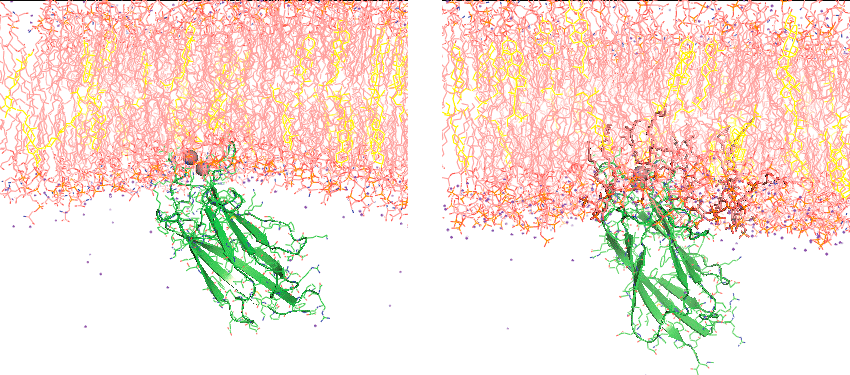
